# Lung basement membranes are compositionally and structurally altered following resolution of acute inflammation

**DOI:** 10.1101/2024.08.19.608567

**Authors:** Oliver Brand, Sara Kirkham, Christopher Jagger, Matiss Ozols, Rachel Lennon, Tracy Hussell, Alexander Eckersley

## Abstract

Identification of pathways preventing recovery from acute respiratory viral infection is under-studied but essential for long-term health. Using unbiased proteomics, we reveal an unexpected persistent reduction in lung basement membrane proteins in mice recovered from influenza infection. Basement membrane provides a critical scaffold for heterogeneous cell types and the proteins they secrete/express at the endothelial and epithelial barrier. Further peptide location fingerprinting analysis shows inherent structure-associated changes within core collagen IV and laminin components, particularly within matrikine-producing regions of collagen IV. Our results imply lingering damage to the basement membrane network despite full recovery from viral infection. Surprisingly, these structure-associated changes in laminin and collagen IV components are also observed in non-infected aged mice indicating that inflammation-driven basement membrane degeneration may contribute to tissue ageing. Interestingly, macrophages in regions deficient in basement membrane express collagen IV and laminin chains. Repair of the basement membrane should therefore be targeted to improve overall lung health.

**Non-technical summary:** Lung virus infection is a constant global threat, despite developments in vaccination and anti-viral treatments. We have a deep understanding of this inflammatory condition, but less is known about the drivers of persistent problems, including fatigue and breathlessness as illustrated by “long COVID”. Here, we reveal a novel finding that a critical structure in the lung (the basement membrane) remains damaged even after the virus and symptoms have cleared. This structure supports a variety of cells that and forms a barrier that lines the airspaces. It also regulates fluid and cell movement into these airspaces. Remarkably, we show that similar persistent changes after virus infection are also evident in aged lungs, which implies that lung complications with age may be due to repeated inflammation. By deciphering the processes causing persistent basement membrane changes, we provide an entirely novel area to target with new medicines to treat complications arising from viral infection.

## Introduction

Resolution of lung inflammation is critical to prevent bystander tissue damage. However, in many cases, resolution is not complete, and the lung accumulates changes over time. These changes are observed in multiple tissue compartments including elevation of extracellular matrix (ECM), formation of ectopic lymphoid aggregates, prolonged disruption to the vasculature and an alteration in the inflammatory tone of the innate immune system [1–4]. Long-term changes are also evident in severe COVID at a time when patients recover and the virus is absent, but a self-sustained local immune response continues to drive tissue remodelling [5].

Matrix remodelling is extensively documented in lung inflammatory disease, though most studies concentrate on matrix products present in excess compared to the healthy state. Deposition extracellular matrix (ECM) affects the stiffness and physiological function of the lung and can alter immune cell retention, trafficking and activation [6–8]. However, there is one common ECM component that remains highly understudied: the basement membrane.

Basement membranes are a form of specialised ECM [9]. They underlie epithelial and endothelial cells forming a structural scaffold to support cells, a template for tissue repair, a reservoir for growth factors, a selectively permeable molecular sieve and an adhesive link between the epithelium/endothelium and the interstitial matrix [10–12]. All basement membranes are composed of two networks of laminin and collagen IV, bridged by linker proteins such as nidogen, perlecan and collagen XVIII [12,13]. These can network further with other assemblies that contribute to more tissue-specific functions [11,12] like the sequestration of growth factors and proteases [14]. Basement membrane collagen IV exists as a trimeric complex, consisting of an N-terminal triple helical coiled coil region connected to a C-terminal, globular NC1 (non-collagenous) domain. Collagen IV trimers associate end to end via this NC1 domain to form hexamers with the α1_2_α2 form being the most abundant of the complexes, present in most basement membranes [9,11,12], and the α3α4α5 form being more tissue-specific and present in lung alveoli [15,16].

An understanding of basement membranes that underlie epithelial cells has lagged behind that of endothelium and there is a paucity of information for the lung. Addressing this gap is important, as defective airway basement membrane is widespread in human diseases, including COVID-19 [17–20]. Damaged basement membrane disrupts tissue structure, barrier and filtration function, allowing serum proteins and red blood cells to leak into airspaces and reducing the ability of epithelial cells to transport proteins of physiological importance [21,22]. Basement membranes also support a variety of epithelial cells expressing or secreting immune modulators that influence airway macrophages, which we know are impaired in the influenza-resolved lung [23–26]. Many cell types, including basal (and suprabasal), club, tuft and epithelial cells, adhere to basement membranes [27] and are required for their epithelial regenerative capacity.

Since basement membranes have a diverse composition of hundreds of proteins which are organised in a hierarchy of macromolecular structures and interactions [11,12], proteomic mass spectrometry (MS) is a useful tool for gauging whether these complex structures return to a pre-inflammation baseline state after infection resolution. Not only has proteomic MS been useful for determining changes in basement membrane composition and relative protein abundance, but novel approaches such as solubility profiling [28] and peptide location fingerprinting (PLF) [29,30] have revealed ageing-related changes in lung ECM architecture and in lung and kidney basement membrane macromolecular structures. Using these techniques, we report that basement membrane composition, protein solubility and hierarchical structure remain altered up to three weeks following acute inflammation. We also reveal specific structure-associated changes within the NC1 domains of four collagen IV chains of the α1_2_α2 and α3α4α5 hexamers, demonstrating network-level disruption to the collagen IV basement membrane meshwork with likely consequences to lung function. Furthermore, we demonstrate that post-inflammatory changes in laminin and collagen IV macromolecular structures are similar to those observed in ageing lung. Finally, we show that lung macrophages localise to areas of damaged basement membranes and their expression of core component proteins significantly increase post-inflammation, which suggests they may harbour the potential for repair.

## Results and Discussion

### Experimental Design

This study was conducted to examine whether tissue matrix returned to baseline following resolution of an acute inflammatory event, with a focus on basement membrane recovery. We used the highly published murine model of influenza virus infection [31,32] administering 5 plaque forming units of influenza A strain APR8 intranasally. This model causes transient weight loss of up to 25 % original body mass with full recovery by day 12. On day 21, lungs were removed and subjected to three rounds of fractionation based on protein solubility **(Figure 1a)**. Equal amounts of protein for each fraction underwent trypsin digestion followed by LC-MS/MS. Agglomerative hierarchical clustering showed a difference in the proteomes recovered from each fraction (indicated by red, yellow and green bars in **Figure 1b**). The greatest number of proteins were identified within the most soluble fraction 1 **(Figure 1c)** which was enriched in intracellular organelle, membrane-bound and cytosolic proteins. The less soluble proteins identified in fraction 2 corresponded to members of the more robust, macromolecular cytoskeleton whereas the least soluble proteins identified in fraction 3 were predominantly ECM associated **(Figure 1d)**.

**Figure 1.**
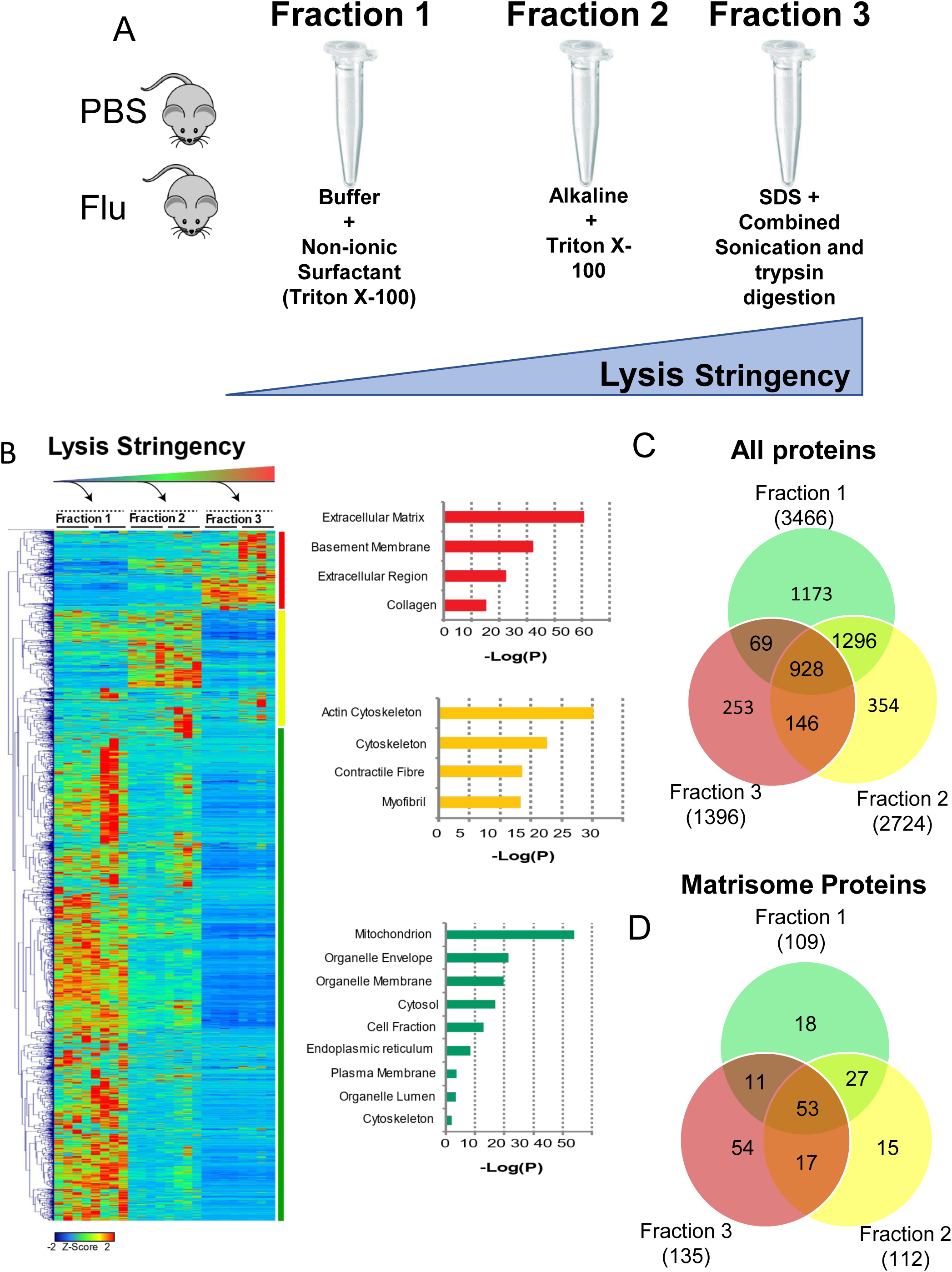
Three fractions generated by increasing lysis stringency yield distinctive proteomes from control and influenza-resolved mouse lungs. Eight-week-old female C57/Bl6 mice (n = 4 per group) were intranasally infected with influenza virus and perfused lungs harvested 21 days later. Minced cell pellets were processed via three rounds of fractionation based on protein solubility and equal amounts of protein for each fraction underwent trypsin digestion followed by LC-MS/MS (A). Agglomerative hierarchical clustering showing a difference in the proteomes recovered from each fraction. Major changes in fraction 1 indicated by the green vertical bar, Fraction 2 the yellow and fraction 3 the red. Bar charts represent a summary of dominant protein classes affected in each fraction (B) Venn diagrams show unique proteins in each fraction and those shared with other fractions in the whole proteome (C) and matrisome (D).

### Basement membrane proteins are reduced in influenza-resolved lung

Using the online resource ‘The Matrisome Project’ [33], fraction 3, containing predominantly basement membrane and ECM proteomes **(Figure 1b,d)**, was interrogated in more depth. The top 15 proteins with the significantly highest down- or up-regulated expressions are shown in Error! Reference source n ot found. and **2** respectively. Of those down-regulated, basement membrane-associated proteins [10] dominated and included collagen IV alpha chains 1-5 (COL4A1-5, ranging from -1.81 to -2.62 fold change), nephronectin (NPNT, -1.67 fold change), laminins alpha 3 (LAMA3, -1.45 fold change), perlecan (HSPG2, -1.60 fold change) and collagen XVIII alpha 1 (COL18A1, -2.31 fold change). Agglomerative hierarchical cluster heat mapping of significantly altered matrisome proteins also showed this reduction in core basement membrane proteins across all influenza-resolved samples (underlined in **Figure 2a**). In particular, the observed reduction of core component laminin (α3, α5 and β2) and collagen IV (α1-5) chains, and stabilising (bridging) components perlecan and collagen XVIII [34], in the post influenza lung, reflects a disruption in basement membrane architecture and integrity. The reduction of nephronectin (NPNT) is interesting as it is a high-affinity ligand for integrin α8β1 and is localised to the alveolar basement membrane. Postnatal deletion of nephronectin results in greater accumulation of airway macrophages in the resolution of LPS-induced inflammation [35]. Pathway enrichment analysis of matrisome-enriched fractions **(Figure 2b)** revealed that altered proteins were not only highly enriched in basement membrane-associated pathways such as laminin interactions, but also in multiple collagen synthesis-associated as well as proteoglycan and cell-integrin interaction pathways. Enrichments plots confirmed a negative correlation for basement membrane and collagen proteins and positive correlation for ECM regulators **(Figure 2c).** Immune fluorescent imaging of pan collagen IV confirmed a reduction of intensity in influenza-resolved (Flu d21) lungs compared to control mice or those at an earlier stage of their infection. There is a particular loss from the bronchioles and in the alveolar wall where the epithelial layer is intact **(Figure 2d)**. Collectively, this represents a global remodelling of matrix composition following influenza resolution.

**Figure 2.**
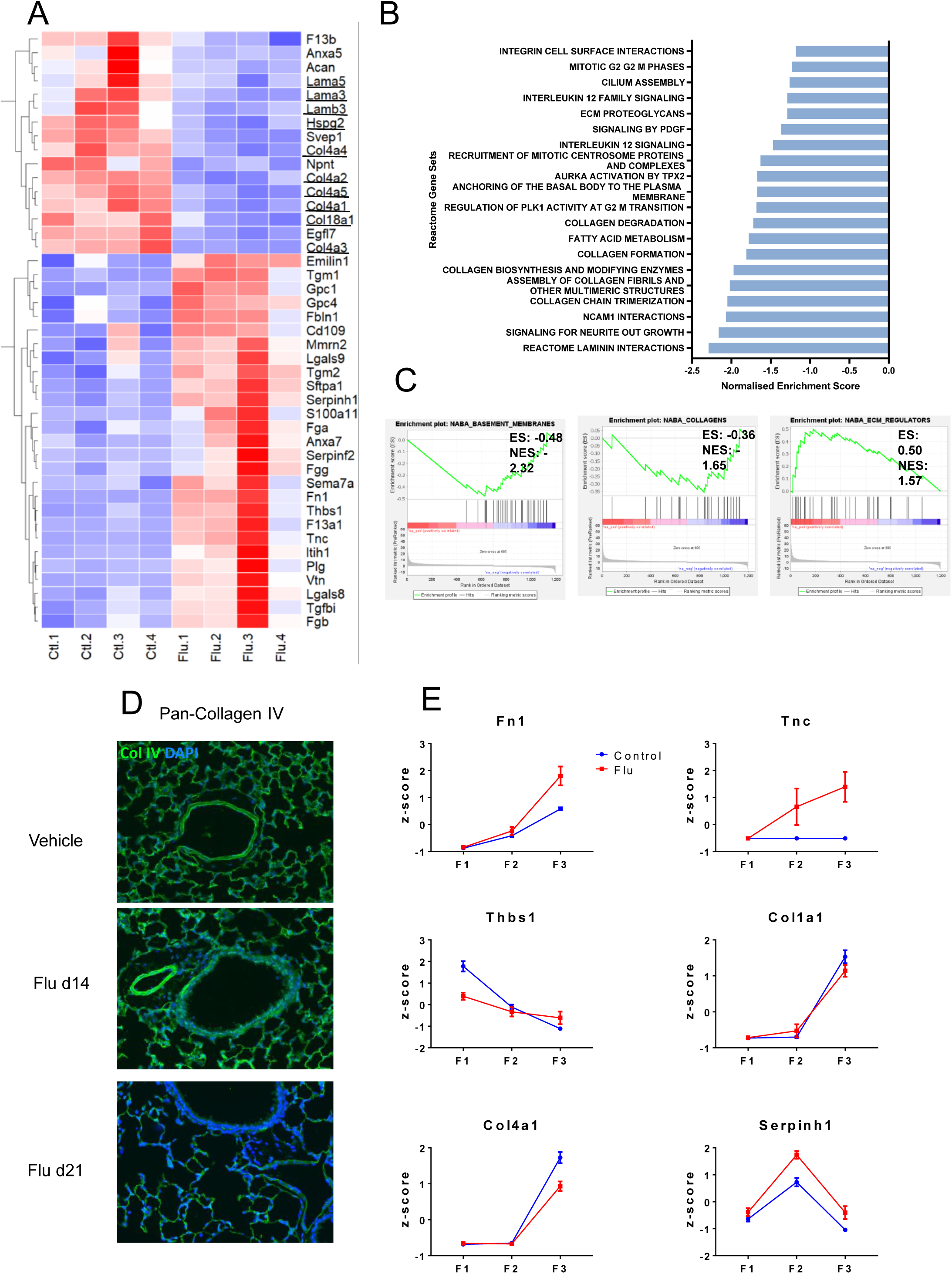
Basement membrane proteins are significantly reduced in mice that have recovered from Influenza infection. Agglomerative hierarchical cluster heatmap of proteins from ECM-enriched fraction three (n = 4 mice) from control (Ctl 1-4) and flu-resolved (Flu 1-4) mice; basement membrane proteins are underlined (A). Pathway enrichment analysis of ECM-enriched fraction three (B). Enrichment plots for up-regulated (red) and downregulated (blue) proteins attributed to basement membranes (left), collagens (middle) and ECM regulation (right) (C). Representative confocal imaging of collagen IV (green) and DAPI (blue) in lungs from control (PBS), and mice 14, and 21 days (n = 6) after infection (D). Profiling of z-scores of indicated proteins across decreasing solubilities in the three extracted fractions (F1-3; n = 5; mean +/- standard deviation [SD]) (E).

A reduction of basement membrane in resolution of influenza virus-induced inflammation could occur for many reasons. Since laminin and collagen IV networks form its core components [36,37], an observed reduction of several of these chains in the post influenza lung has significant implication, as does the reduction of perlecan, collagen XVIII and heparin sulphate proteoglycan that stabilise the basement membrane [34]. Influenza virus infection can induce autoantibodies to exposed collagen IV and is associated with development of both anti-glomerular basement membrane disease (Goodpasture syndrome) and alveolar damage [38,39]. Cellular transmigration can occur through regions of low basement membrane protein presence and neutrophils, in particular, migrate through these regions by producing proteases [40]. However, most studies have focussed on endothelial basement membrane and transmigration through the often-fused combination of endothelium and epithelium of the lower airways remains obscure. Matrix metalloproteinases (MMP), particularly MMP-2 and -9 are associated with basement membrane degradation and are induced in hypoxia and inflammation [41]. During influenza virus infection, MMP9 is observed in airway epithelial cells and infiltrating immune cells and MMP9 knockouts (-/-) develop less severe disease to influenza virus infection [42]. We previously reported that inhibition of MMP9 prevents the development of influenza-induced bacterial superinfection in the lung [43]. Though basement membrane integrity was not examined at that time, it is widely known that prior influenza infection facilitates bacterial entry into the lung tissue and blood stream; an effect not observed in bacterial infection alone.

The last 184 amino acid residues of collagen XVIII α1 (which was significantly downregulated in influenza-resolved lung **[Table 1**, **Figure 2a]**) contains the matrikine endostatin (20 kDa) that is released along with endostatin-like fragments by the actions of elastase or cathepsin L [44,45]. As well as integrins, endostatin also interacts with the glypican (GPC) cell surface receptors [46], two of which (GPCs 1 and 4) were up-regulated (GPC1 significantly) in the influenza-resolved lung **(Figure 2a).** Endostatin is a potent inhibitor of endothelial cell proliferation and migration, and as such has been examined for its anti-angiogenic properties in cancer and as a biomarker for chronic obstructive pulmonary disease (COPD) [47]. However, endostatin is now known to possess many other functions including in apoptosis [48]. Unfortunately, we did not detect enough collagen XVIII peptides to quantify changes in the NC1 domain associated to endostatin, however this merits further investigation.

**Table 1.** Most highly decreased proteins in the matrisome enriched fraction 3 in mouse lung at day 21 after influenza virus infection.

| Gene Symbol | Fold Change | p-value | Comments | Reference |
| --- | --- | --- | --- | --- |
| **Svep1** | -4.30 | 0.0002 | Sushi, Von Willebrand Factor Type A, EGF And Pentraxin Domain Containing 1<br>Potential biomarker for lung cancer with metastatic pleural effusion | [82] |
| **Anxa5** | -4.06 | 0.0144 | Annexin A5<br>Increased in lung cancer and COPD BALF<br>Binds to phosphatidylserine | [83,84] |
| **Egfl7** | -3.51 | 0.0002 | EGF-like domain-containing protein 7<br>Has sequence for miR-126 and miR-126 in intron 9.<br>Promotes cells invasion and angiogenesis in pancreatic carcinoma | [85,86] |
| **Col4a5** | -2.62 | 0.0006 | Collagen Type IV alpha 5<br>Part of basement membrane (minor Col IV).<br>Required for lung cancer progression | [87] |
| **Acan** | -2.61 | 0.0189 | Aggrecan<br>Involved in wound healing/repair in skin (high in granular phase, then decreased) and cartilage | [88] |
| **Col4a3** | -2.59 | 2.59E-05 | Collagen Type IV alpha 3<br>Part of basement membrane. Has suppressed expression in COPD. | [89] |
| **Col18a1** | -2.31 | 0.0028 | Collagen Type XVIII alpha 1<br>Part of vascular/epithelial BMs. Associated with poorer prognosis in non-small cell lung carcinoma | [90] |
| **Col4a4** | -2.08 | 0.0007 | Collagen Type IV alpha 4<br>In alveolar BM of normal lung. Get gradual loss in bronchioloalveolar carcinoma. | [15] |
| **Col4a1** | -1.93 | 0.0004 | Collagen Type IV alpha 1<br>Deposited in fibroblastic foci in usual interstitial pneumonia and inhibit fibroblast migration.<br>Stimulated by TGF-β | [91] |
| **F13b** | -1.91 | 0.0128 | Coagulation Factor XIII, B Polypeptide<br>Linked to cystic fibrosis | [92] |
| **Col4a2** | -1.81 | 0.0001 | Collagen Type IV alpha 2<br>Deposited in fibroblastic foci in usual interstitial pneumonia and inhibit fibroblast migration.<br>Stimulated by TGF-β | [91] |
| **Npnt** | -1.67 | 0.0034 | Nephronectin | [93,94] |
| | | | BM component. Downregulated by TGF-β. Loss promotes metastasis in malignant melanoma. | |
| **Hspg2** | -1.60 | 0.0031 | Heparan Sulfate Proteoglycan 2 (perlecan).<br>BM component. Upregulated by TGF-β in airway smooth muscle cells from COPD. | [95] |
| **Lama3** | -1.45 | 0.0074 | Laminin alpha 3<br>BM component. Reduced in fibrotic lung regions | [96] |

### Matrix adherence glycoprotein abundance and solubility is altered in influenza resolved lung

Interestingly, the fibrosis-linked [49], cell-matrix adhering glycoproteins fibronectin (FN1; 8.39 fold change), vitronectin (VTN; 3.07 fold change) and tenascin C (TNC; 8.65 fold change) were all significantly higher in influenza-resolved lung compared to control **(Table 2**, **Figure 2a)**.Due to these links to fibrosis, which can decrease the solubility of tissues, we next used quantitative detergent solubility profiling [28] to interrogate whether the solubility profiles across increasing detergent stringencies of key matrix proteins differed between influenza-resolved and control lung **(Figure 2e)**. The glycoproteins fibronectin, tenascin C, and thrombospondin-1, all exhibited different solubility profiles between influenza-resolved lung and control samples whereas the profiles of collagens I and IV were similar when compared with the control. Since these glycoproteins exist in pericellular spaces where they mediate cell-matrix interactions [50], this indicates an alteration in local pericellular matrix architecture following influenza resolution and could reflect a change in local integrity and cell-matrix adhesion.

**Table 2.**
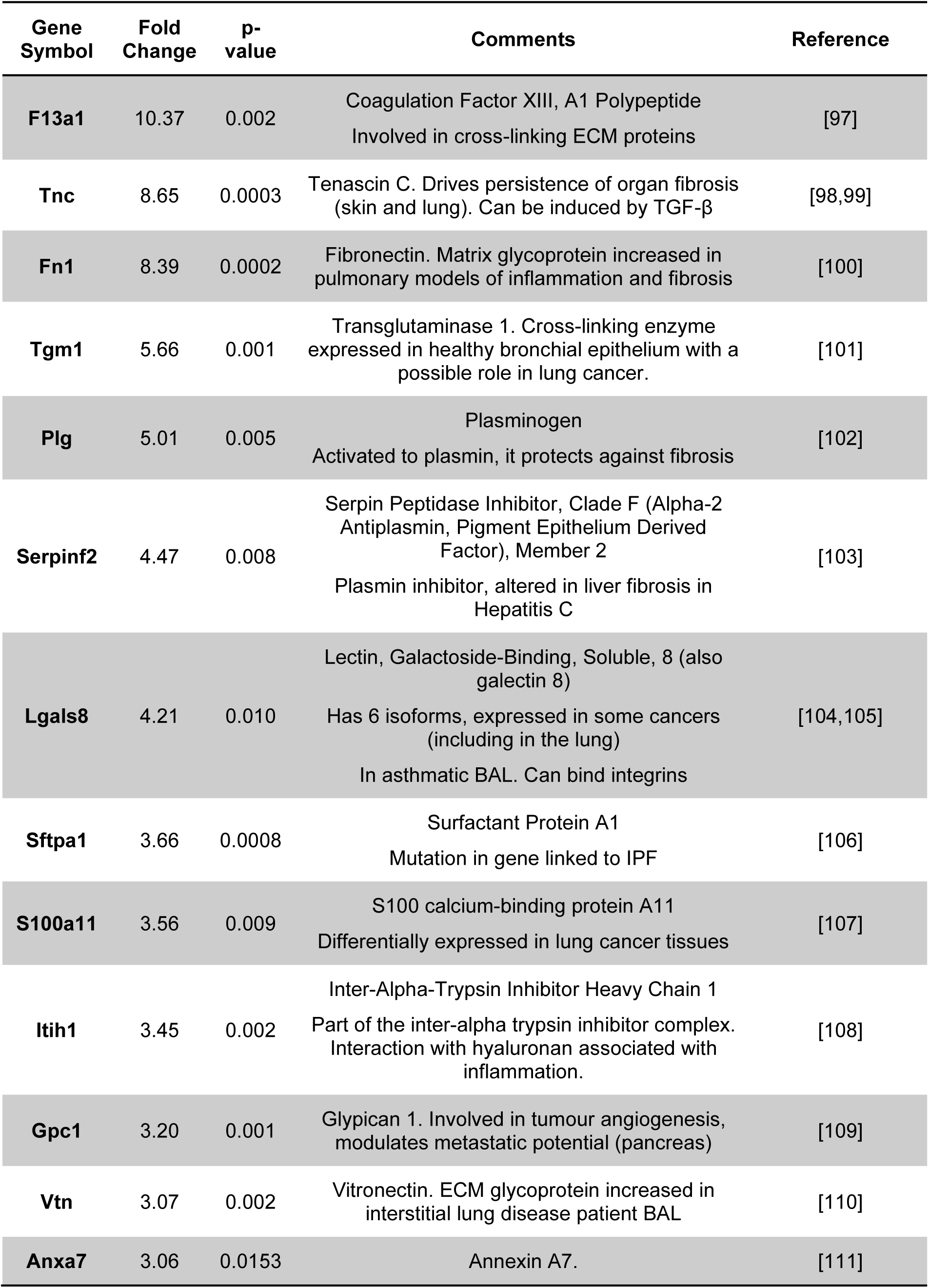
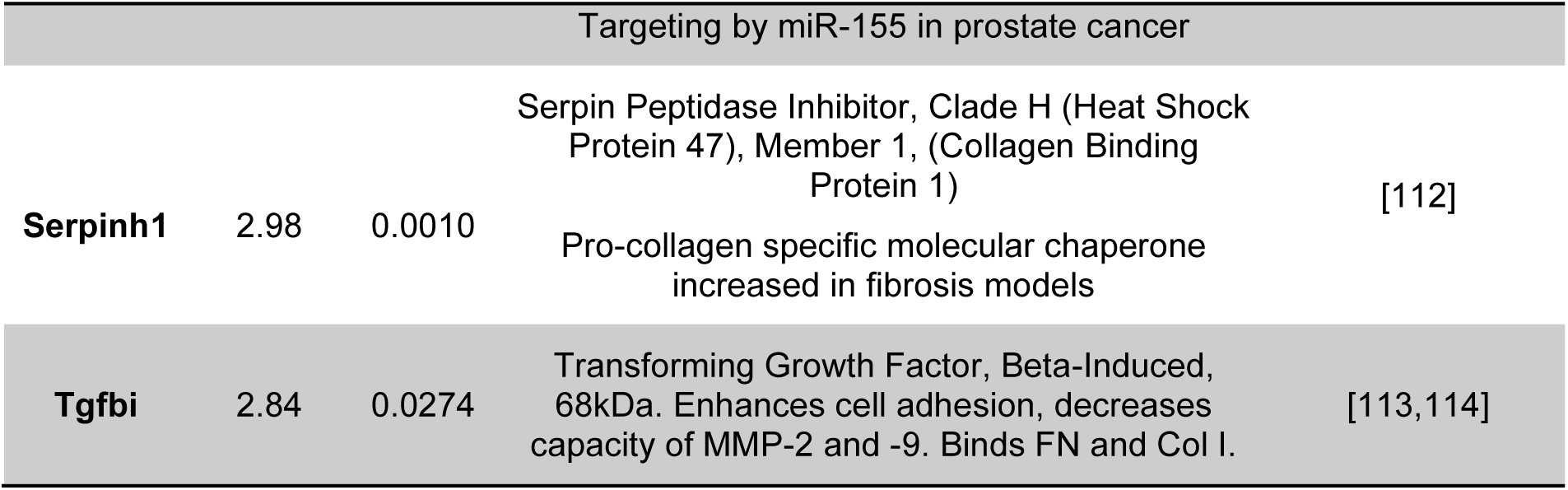
Most highly increased proteins in the matrisome enriched fraction 3 in mouse lung at day 21 after influenza virus infection.

### Basement membrane proteins harbour structure-associated differences in Influenza-resolved lung compared to control

To investigate whether basement membrane proteins in influenza-resolved lung harboured structure-associated differences that may reflect changes in their macromolecular organisation and interactions, LC-MS/MS datasets of the ECM-enriched third fractions **(Figure 1b,d)** were analysed by peptide location fingerprinting (PLF).

Proteotypic peptides **(Table S1)** were identified by MS2 ion searches. Principal component analysis of peptide spectral counts demonstrated good data separation between groups, with control samples (PBS) clustering along PC1 (indicative of similar compositions), in contrast to Influenza-resolved samples (Flu), which were comparatively more variable **(Figure S1)**.

A total of 149 proteins were identified by PLF with significant differences in peptide yield along their structures **(Table S2)**. Classification analysis **(Figure 3a)** revealed ECM proteins as one of the top five classes affected, constituting 20% of these shortlisted biomarker candidates, with 13% being basement membrane associated. These included core basement membrane component proteins such as four collagen IV alpha (α) chains, four laminin chains and two nidogens, as well as basement membrane co-localising proteins such as collagens VI and XII, fibronectin, the matricellular protein periostin, and the proteoglycan versican. Holistically, this indicates that basement membranes in influenza-resolved lungs are structurally distinct (on a molecular level) from those present prior. In addition to ECM proteins, cytoskeletal proteins (e.g. several keratins and myosins), protein modifying enzymes (e.g. matrix metalloproteinase 12 and aminopeptidase N), and metabolite interconversion enzymes (e.g. peroxiredoxin-1) were also in the top classes structurally affected, indicating a change in wider proteostasis in Influenza-resolved lung.

**Figure 3.**
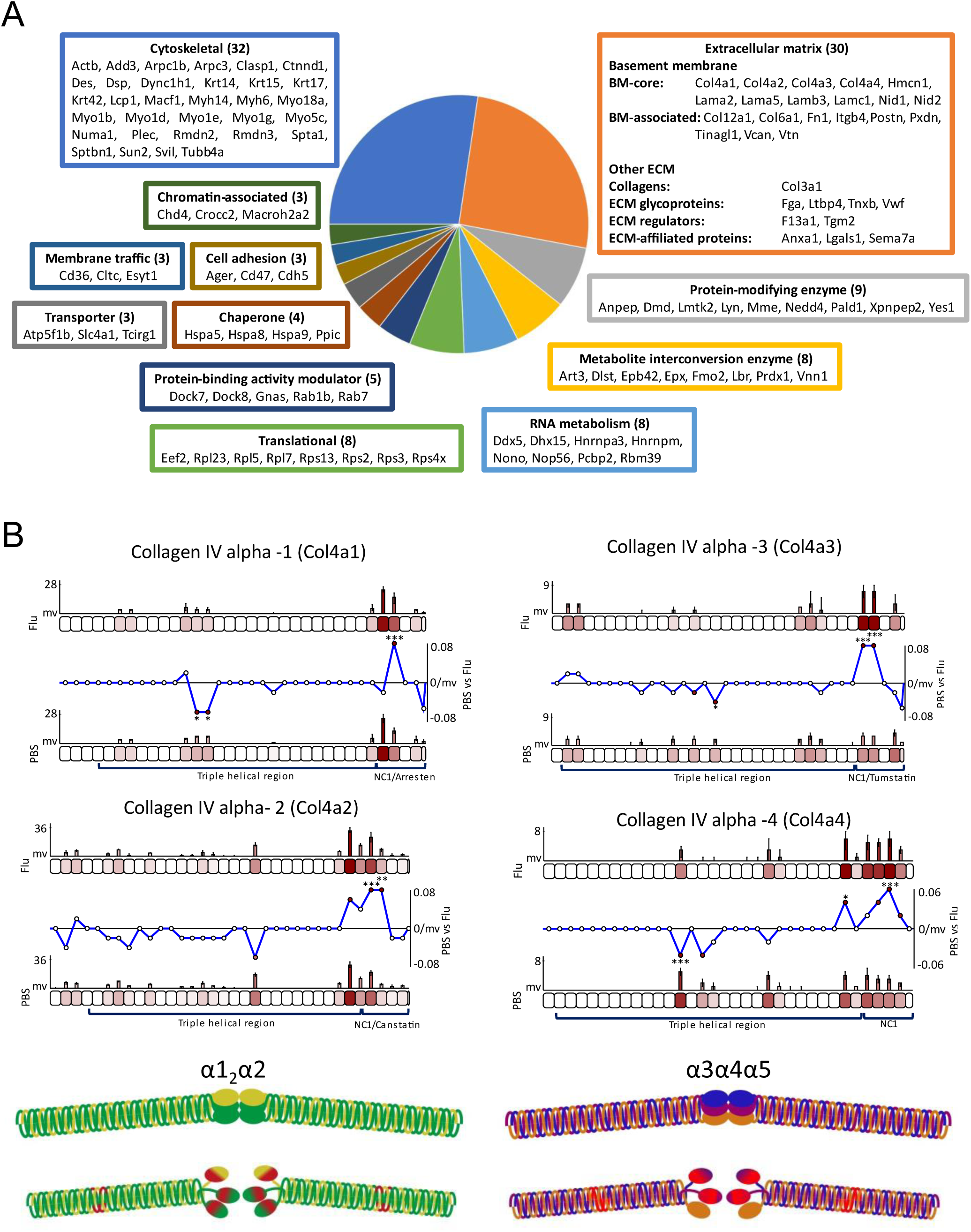
The Collagen IV network from Influenza-resolved lung is structurally distinct to those present in non-infected mice. Classification analysis (using BMbase [11] for basement membrane proteins, MatrisomeDB [115] for ECM and PANTHER [116] for all others) of the 149 proteins identified by PLF with structure-associated differences revealed basement membrane components as one of the main categories affected (A). Classes and their constituent proteins (total number of proteins per class indicated in brackets) are displayed in coloured frames corresponding to their respective pie chart slice. Only classes with 3 proteins or more are shown. Collagen IV α chains 1 – 4 from Influenza-resolved lung exhibit markedly conserved differences in peptide yield along their structures compared to control (B). During PLF analysis [29,51,81], proteins were divided in 50 amino acid (aa)-sized segments and LC-MS/MS-identified, proteotypic peptides were mapped, quantified (bar graphs = average peptide spectral counts, median normalised based on protein-specific total spectral counts per segment, error bars = SD, missing values are indicated at y = 0 [mv]; line graphs = average normalised spectral count in PBS controls subtracted from Influenza-resolved and divided by the segment length [50 aa], to reveal peptide yield differences along structures between the two groups) and statistically tested (unpaired Bonferroni-corrected, repeated measures ANOVA; *, p ≤ 0.05; **, p ≤ 0.01; ***, p ≤ 0.001) between Influenza-resolved and control (PBS) samples. Functional domains and regions (sourced from Uniprot) are also indicated.

### The collagen IV network from Influenza-resolved lung is structurally distinct to those present in non-infected

To explore the potential functional consequences of the significant differences in peptide yields identified along the structures of basement membrane proteins, these were correlated to functional and conserved structural regions, as previously done [29,30,51]. Additionally, proteins which formed complexes and polymeric networks (e.g. laminin and collagen α chains) were also examined for changes that might exist on a quaternary level [30].

In particular, four collagen IV α chains (α1–α4) were all found to exhibit significantly higher peptide yields specifically within their C-terminal non-collagenous (NC) 1 domains in Influenza-resolved samples compared to control **(Figure 3b)**. Conversely, lower peptide yields were also consistently observed within the triple helical regions of all four of these collagen IV α chains (significant within the central portions of this region for all chains except α2).

The peptide yield difference patterns that appear markedly conserved between collagen IV α1, α2, α3 and α4 chains can be better interpreted by examining the complexes and networks they form. From the six types of collagen IV α chains available (α1– α6), only three combinations of triple helical protomers exist in tissue: α1_2_α2, α3α4α5 and α5_2_α6, all of which are capable of binding end-to-end to form hexamers with protomers of the same kind [52]. The α1_2_α2 hexamer is the most abundant of the complexes and present in all basement membranes [9], whereas the α3α4α5 and α5α5α6 hexamers are more tissue-specific, with the α3α4α5 complex being present in lung alveoli [15,16]. When visualised in context of their respective α1_2_α2 and α3α4α5 hexamers, we observed that the significant differences in proteotypic peptide yields across α1–α4 from Influenza-resolved lung overlapped in similar locations found within their hexamer’s globular NC1 domains and triple helical regions (**Figure 3b**, coloured red in schematic models). The fact that the same, conserved differences were observed across four α chains that form two distinct hexamers suggests that these changes may permeate a large proportion of the lung collagen IV basement membrane meshwork with likely consequences to lung function. Furthermore, the largest significant differences in post-influenza collagen IV α1–α4 were consistently seen within their NC1 domains which are crucial for the formation of hexamers through an NC1-NC1 aminoterminal interaction [52,53]. Changes within this domain are therefore also suggestive of a network-level change in collagen IV interaction following infection resolution. Importantly, these observations, in addition to the decrease of core components in influenza-resolved lung, **(Table 1**, **Figure 2)**, suggest that basement membranes that are laid down following Influenza resolution may be structurally and compositionally distinct **(Figure 3b)** from those laid down during growth, or development.

The NC1 domains of collagen IV α1-α4 are also susceptible to fragmentation by MMPs which produces four distinct cell signalling matrikines: arresten, canstatin, tumstatin and tetrastatin respectively [54]. These bioactive fragments can exert a number of biological effects in tissue via interactions with cell integrins, including inhibition of angiogenesis (arrestin and canstatin) and epithelial and fibroblast cell proliferation and migration (canstatin, tumstatin and tetrastatin) [54]. It is interesting to speculate therefore whether the differences in peptide yields observed within the NC1 domains may relate to changes in the production of these fragments in Influenza-resolved lung, which merits further investigation.

### Basement membrane laminin and collagen IV chains from Influenza-resolved lung exhibit similar structure-associated differences to those in ageing

As in all organs, the lung ages leading to a progressive decrease in function and capacity over time. In contrast to intracellular proteins, many components of the extracellular matrix are long-lived [55] (e.g. human elastic fibre [56] and collagen I [57] turnovers span decades), including basement membrane proteins [58]. As such, matrix proteins are vulnerable to the accumulation of structural damage in ageing, such as from oxidation and fragmentation by chronic exposure to reactive oxygen species (ROS) and proteases respectively [59–61], which may impact on their function. Inflammatory processes are known to remodel and degrade long-lived matrix components such as those in basement membranes [62], which without fast and effective mechanisms of repair, may contribute to the structural damage identified in ageing lung. In this way, we propose that inflammation-driven degeneration of matrix may contribute to and potentially accelerate ageing in tissues such as lung.

To investigate whether matrix components from influenza-resolved lung might exhibit similar structure-associated changes to those in ageing, peptide yield difference patterns from biomarker candidates identified here **(Figure 3a)** were compared to those also identified by PLF in a previously published comparison between young (3 months old) and aged (24 months old) mouse lung [29]. In that study, a total of 140 proteins were identified by PLF from ECM-enriched bulk lung samples with structure-associated differences in ageing [29]. Of these, 26 were also identified here with post Influenza-specific differences in structure **(Figure 4a).** Remarkably a large proportion these proteins (11, 42%) were basement membrane affiliated, which suggests that basement membranes from Influenza-resolved lung may exhibit an aged-like phenotype.

**Figure 4.**
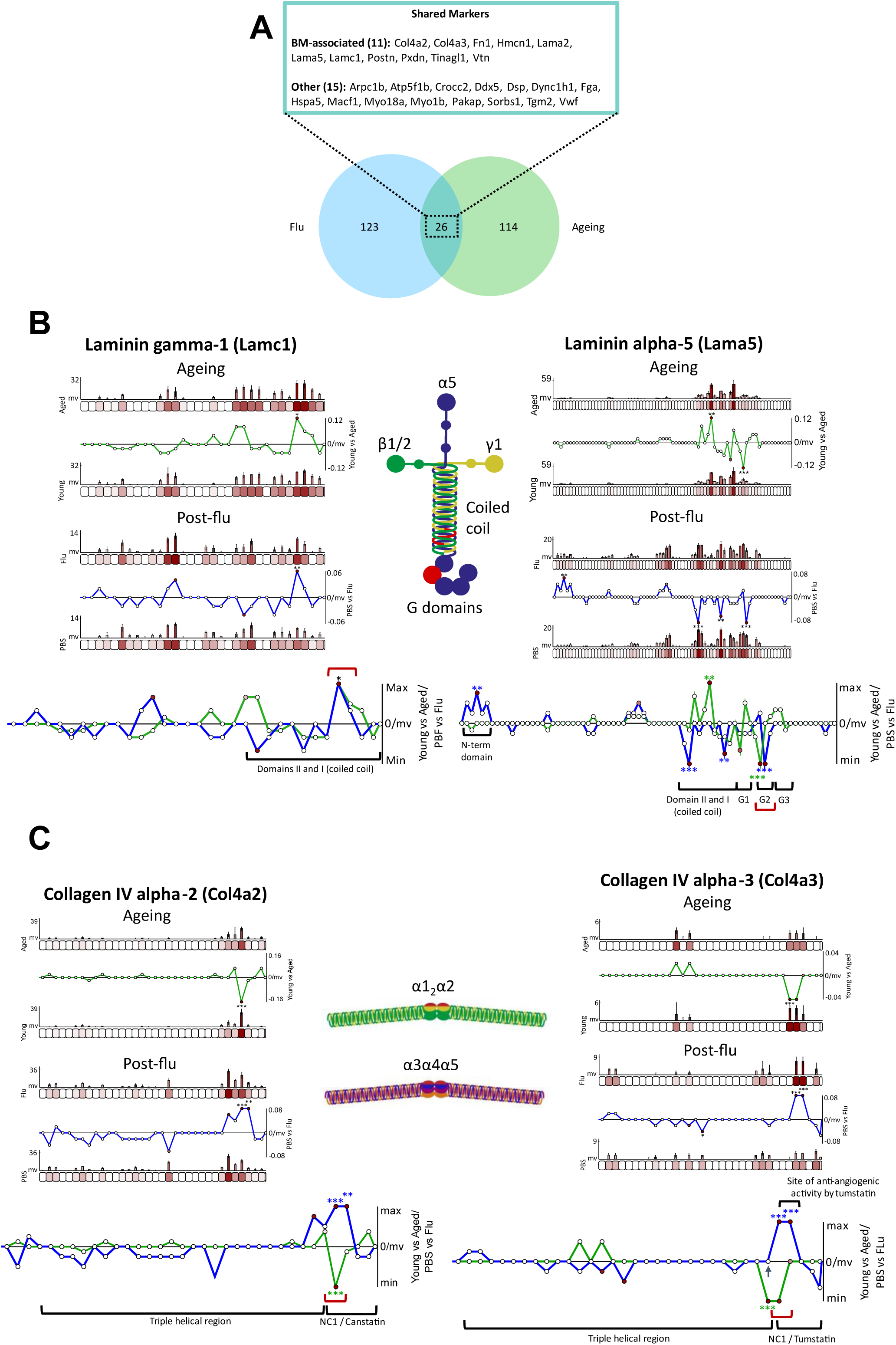
Post-Influenza changes in basement membrane proteins match changes observed in aged mice. Twenty-six proteins were identified by PLF with significant structure-associated differences in both aged (compared to young) and Influenza-resolved (compared to control) mouse lung (A). Of these, 42% were BM-associated (classified using BMbase [11]), including multiple collagen IV and laminin chains. Laminin α5 and γ1 (B) and collagen IV alpha-2 and -3 (C) chains exhibited regional structure-associated differences which were conserved between mouse lung ageing and post-flu resolution. For description of graphs produced by PLF analysis, please see Figure 3. The composite line graphs consist of both aged vs young (green) and post-flu vs PBS control (blue) lines, normalised and overlaid to highlight protein regions with conserved differences between lung ageing and resolved flu (red brackets).

Of the basement membrane proteins which were found to have structure-associated differences in both influenza-resolved and aged lung, laminins α5 and γ1 and collagen IV α2 and α3 chains all exhibited similar regional differences in peptide yields. Both laminin chains exhibited differences within the globular G2 domain for α5 and in the same segment within the coiled coil region of γ1, which were conserved between aged and Influenza-resolved lung (**Figure 4b**, red brackets). Similarly, collagen IV α2 and α3 chains both harboured significant differences in peptide yield within a similar protein region, corresponding to the C-terminal NC1 domains (**Figure 4c**, red brackets), however, these were higher in Influenza-resolved and lower in aged.

Laminins α5 and γ1 complex to form two distinct isoforms of α5β1γ1 (511) and α5β2γ1 (521) configurations in lung [63]. Within these complexes, the α chain’s G2 domain and the C-terminal region of the γ chain’s coiled coil are both crucial for integrin-mediated cell interaction [64]. The conserved, structural differences observed within these regions **(Figure 4b)** may therefore reflect a dynamic shift in cell-laminin interaction state or a perturbation in the ability of these laminins to bind cells; both phenotypes being potentially shared between aged and influenza-resolved lung.

As discussed previously, the α2 and α3 chains of collagen IV each form two distinct α1_2_α2 and α3α4α5 hexamers [52,53]. Although significant peptide yield differences are reversed in these two chains between aged (lower than young) and Influenza-resolved (higher than control) lung, the same regions corresponding to the NC1 domains of the collagen IV hexamers were affected **(Figure 4c)**, suggesting that this macromolecular region is susceptible to lasting changes both in both ageing and post infection. With the NC1 domain being important for the complexing and networking of collagen IV [52,53], changes within both α1_2_α2 and α3α4α5 hexamers suggest a basement membrane-wide effect which occurs in both ageing and after recovery from acute inflammation. As mentioned previously, the MMP-mediated fragmentation of the α2 and α3 NC1 domains (**Figure 4c** black arrow) releases the functional canstatin and tumstatin matrikines [54], therefore changes within this region could also reflect differences in the presence of these fragments in both aged and Influenza-resolved lung.

Changes within the same macromolecular regions of laminins 521/511 and collagen IV in both ageing and Influenza-resolved lung suggest that effects of acute inflammation may cause lasting damage to basement membranes on a molecular level which remain present in later life.

### Airway macrophages have the potential to repair basement membrane

It remains unclear which cells contribute to basement membrane repair in tissue. *In vitro*, the mouse lung epithelial (MLE)-15 cell line, which is similar to alveolar epithelial type II cells, produces laminin α5, and trace amounts of α3, β1 and γ1 chains, collagen IV and perlecan [63]. When grown on fibroblast-embedded collagen gels, MLE-15 cells assemble a basement membrane-like layer. Similarly, immortalized alveolar type II epithelial (SV40-T2) cells form continuous, thin layers of *lamina densa* when cultured on collagen fibrils with supplemental transforming growth factor (TGF) β1. Integration of laminin, type IV collagen, perlecan, and nidogen was also confirmed as part of this underlying layer [65,66]. We also observe pan-collagen IV staining in alveolar epithelial cells and the larger airways that is reduced in influenza-resolved mice (**Figure 2d and 5a**). More surprising, however, was the presence of strong staining for collagen IV in macrophages clustered in regions of epithelial damage **(Figure 5a)**. The idea of macrophage production of collagen IV is complicated by the fact that these cells are responsible for its clearance. Uptake of collagen into macrophages occurs by phagocytosis of intact fibrils and by micropinocytosis or receptor-mediated endocytosis of cleaved fragments [67]. Subsequent intracellular degradation involves lysosome fusion and the activity of cathepsins [68].

**Figure 5.**
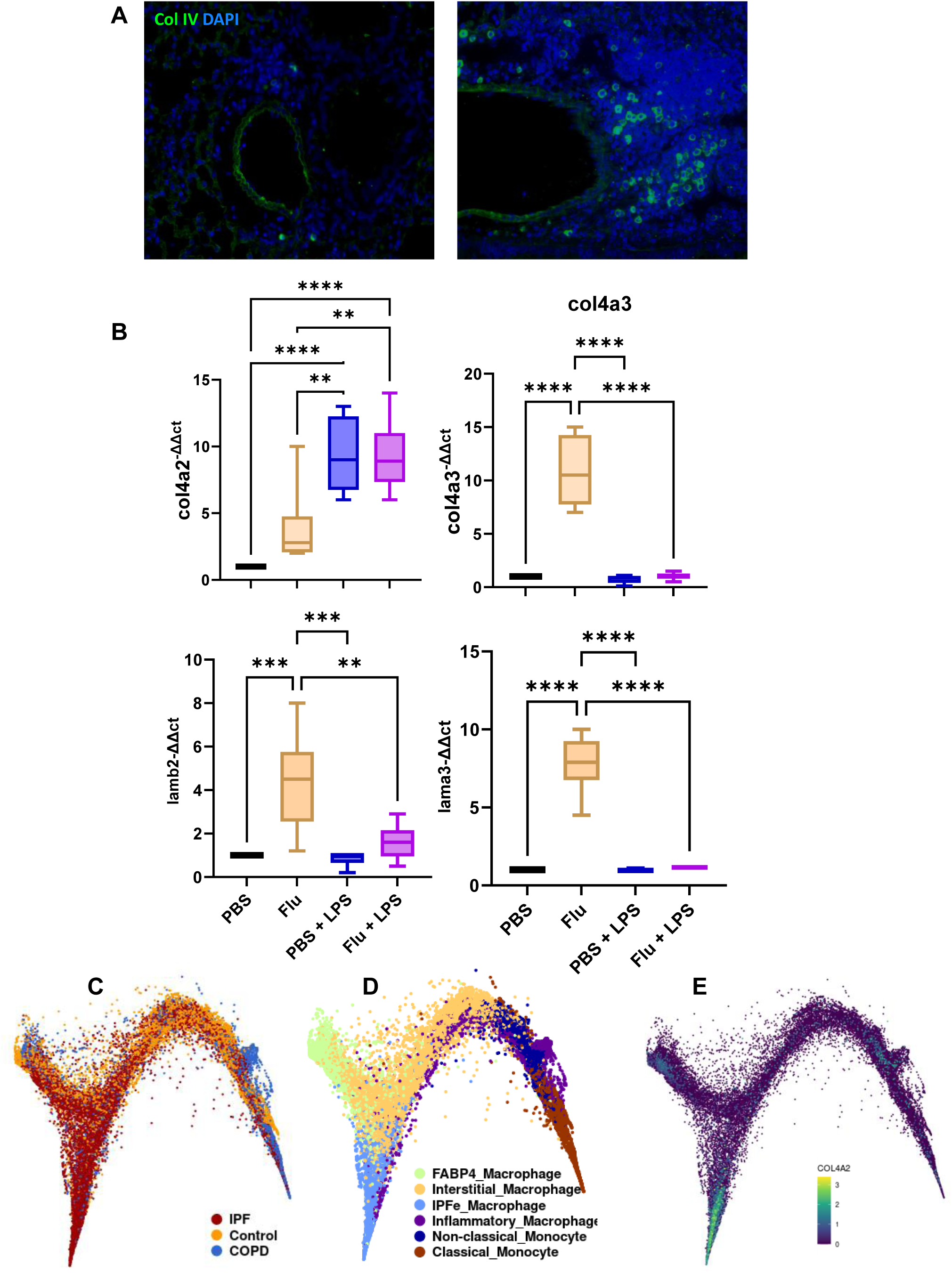
Macrophage production of basement membrane components. Immunofluorescence images of lung sections from mice administered with PBS (left) or influenza virus (right) showing DAPI in blue and pan-collagen IV in green (Representative of n = 6; magnification x40) (A). RT-PCR analysis of FACS purified whole lung macrophages (using CD206 [Mannose receptor] and CD169 [Siglec-1]) from mice given PBS or influenza (Flu) 21 days earlier. Macrophages from these groups were also stimulated *in vitro* with 10 μg LPS for 24 hours and RT-PCR analysis performed for the indicated mRNA transcripts. Results shown as a mean + min-max box and whisker and significance calculated using an unpaired Bonferroni-corrected, repeated measures ANOVA (**, p ≤ 0.01; ***, p ≤ 0.001; ****, p ≤ 0.0001) (B). Reanalysis of an online dataset of macrophages from idiopathic pulmonary fibrosis [48] showing their distribution by UMAP based on disease (C) macrophage subtype (D) and Col4a2 (E).

As intracellular protein presence could reflect uptake, we examined mRNA transcripts of basement membrane components by RT-PCR in macrophages under different stimulatory conditions. Cultured macrophages isolated from influenza-resolved (Flu) lungs expressed higher levels of collagen IV α3 and laminin α3 and β2 transcripts compared to macrophages from negative controls (PBS) **(Figure 5b)**. Additional *in vitro* stimulation of PBS control and influenza resolved lung macrophages with LPS enhanced col4a2 but not other basement membrane core proteins. This suggest that different macrophage stimuli induce different basement membrane components: influenza, Col4a3, Lamb2 and Lama3, and LPS Col4a2.

The observation that macrophages localise to basement membrane regions of influenza-resolved lung and express higher levels of basement membrane core proteins (**Figure 5b**), suggests that they may harbour the potential for repair. Expression of basement membrane components by certain subspecies of macrophages also occurs in idiopathic pulmonary fibrosis (IPF). Uniform manifold approximation and projection (UMAP) plots (**Figures 5c**), generated using the IPF cell atlas (https://p2med.shinyapps.io/IPFCellAtlas/_w_5a85c8d9/#tab-6473-3; last accessed 8^th^ August 2024), from recent single-cell RNA-seq by Adams *et al.* [69] indicate a subpopulation of IPFe-expanded macrophages **(Figure 5d)** that uniquely express Col4a2 **(Figure 5e)**, when compared to other subtypes. Similarly, recent proteomic analysis by Oates *et al*. also shows that peripheral blood macrophage subtypes, cultured for seven days, differentially express collagen IV α2 and α5 chains, with the M2c subtype producing higher levels than control (M0) macrophages and M1 subtypes producing less [70].

The concept of whether macrophages, which are known for their role in basement membrane damage during transmigration, also exist as highly mobile sentinels of repair is an interesting one, and merits future consideration. This may even be evolutionary conserved: Matsubayashi *et al* [71] showed that migrating macrophages (haemocytes) in *Drosophila* embryos are not only the largest producers of basement membrane, but that this role is critical to embryonic development.

## Conclusion

Through proteomic and PLF analysis, we not only showed a reduction in lung basement membrane proteins, following influenza-resolution, but also identified inherent structure-associated changes within the functional domains of core collagen IV and laminin components, reflecting lingering damage to basement membrane network and architecture despite full recovery. Although the mechanism impeding basement membrane repair has not yet been elucidated, the observation that influenza-resolved lung yields significantly more peptides within matrikine-producing regions of collagen IV merits further investigation. This may represent either a disruption in local homeostasis or serve as a mechanism of cell recruitment and repair. We also observed that inflammatory processes targeted the same protein regions of laminin and collagen IV chains as seen in ageing lung, providing evidence that that inflammation-driven ECM degeneration may accelerate or contribute to tissue ageing. Interestingly, we found that macrophages sourced from influenza-resolved lung readily express collagen IV and laminin chains, revealing a potential role in basement membrane repair for these highly motile cells and providing an exciting new avenue of regenerative research.

## Materials and Methods

### Influenza infection and tissue processing

All experiments on animals were undertaken in accordance with the UK Home Office Animals (Scientific Procedures) Act of 1986 and European Directive 2010/63/EU. Mice were housed in a 12-h light/dark cycle at 21°C with free access to food and water. Female mice (n = 4 per group, 12 weeks old) were intranasally infected with 5 plaque-forming units (pfu) of influenza A virus, Puerto Rico/8/34 (H1N1) in 30 µl PBS and weight recorded daily. Mice were culled at d 21 after infection, perfused with PBS + 1 mM EDTA via the right ventricle to remove blood and lungs snap frozen in liquid nitrogen and stored at – 80 °C. In some experiments lungs were inflation-fixed in formalin prior to paraffin embedding. 5 μm slices were examined for collagen IV staining by immunofluorescence and intensities compared with ImageJ. Some macrophages isolated from Influenza-resolved or PBS control mice were incubated with 10 μg LPS for 24 hours prior to analysis of basement membrane proteins by qPCR.

### Protein extraction for mass spectrometry

Protein extraction was based on previously published methods [72]. Whole lung lobes were weighed, minced, suspended in ice cold TB 5:1 buffer volume to tissue weight (10mM Tris, 150mM NaCl, 25mM EDTA, 1% (v/v) Triton X-100 with 25 μg/ml Leupeptin, 25 μg/ml Aprotinin and 0.5 mM AEBSF), sheared through a 21G needle and left on a rotor overnight at 4°C. Samples were centrifuged at 14,000 g, 10 min at 4°C and the supernatant collected and frozen (Fraction 1). For fraction 2 the pellet was then re-suspended in ice cold EB (20mM NH4OH, 0.5% (v/v) Trition-X100 in PBS), sheared and left on a rotor for 1 hour at 4°C. Samples were centrifuged and the supernatant stored (Fraction 2). The pellet was then re-suspended in PBS with DNase (25μg/mL, 18047019, ThermoFisher) and RNase (25μg/mL, EN0531, ThermoFisher), rotated for 30 min at RT and then centrifuged and the supernatant discarded (unused fraction). Pellets were re-suspended in guanidine solution (4M Guanidine hydrochloride, 50mM sodium acetate, 25mM EDTA with protease inhibitors, pH 5.8) and rotated for 24-48 hours at RT until samples were fully dissolved. Proteins were precipitated with 10:1 volume absolute ethanol at -20°C overnight, centrifuged at 16,000g for 45 min, the pellets washed with 90% ethanol, then re-suspended in 5x sample buffer (7% SDS, 30% glycerol, 0.2M Tris-HCl, 0.01% bromophenol blue, 10% β-mercaptoethanol, pH 6.8) (Fraction 3).

All protein samples were resolved by SDS-PAGE and visualized by Coomassie InstantBlue (ISB1L, Expedeon, Cambridgeshire, UK) staining. Gel tops were subjected to in-gel trypsin digestion. Gel pieces were de-stained three times with 50% Acetonitrile and 50% ammonium bicarbonate solution for 30 minutes, dehydrated by immersing in acetonitrile followed by vacuum centrifugation for 30 minutes. Subsequently, gel pieces were reduced in 10 mM dithiothreitol (DTT), alkylated in 55 mM iodoacetamide, and washed alternatively with ammonium bicarbonate and acetonitrile. Gel pieces were then dehydrated and digested with sequencing grade trypsin (12.5ng/μL). Peptides from gel slices were collected in one wash of 99.8% (v/v) acetonitrile, and 0.2% (v/v) formic acid and one wash of 50% (v/v) acetonitrile and 0.1% (v/v) formic acid. Peptides were desiccated in a vacuum centrifuge and resuspended in 50 μl of 5% (v/v) acetonitrile and 0.1% (v/v) formic acid.

After desalting all samples were run on a nanoACQUITY UltraPerformance liquid chromatography system (Waters, Elstree, UK) coupled offline to an Orbitrap Elite analyser (ThermoFisher). Peptides were separated on a bridged ethyl hybrid C18 analytical column (250 mm length, 75 μm inner diameter, 1.7 μm particle size, 130 Å pore size; Waters) using a 45-min linear gradient from 1% to 25% (v/v) acetonitrile in 0.1% (v/v) formic acid at a flow rate of 200 nl/min. Peptides were automatically selected for fragmentation by data-dependent analysis.

### Proteomic mass spectrometry data analysis

Tandem mass spectra were extracted using extract_msn (ThermoFisher) and loaded into Mascot Daemon (version 2.5.1; Matrix Science, London, UK). Peak list files were searched against a modified version of the Uniprot mouse database (version 3.70; release date January 2015), containing ten additional contaminant and reagent sequences of non-mouse origin, using Mascot (v2.5.1, Matrix Science, MA, USA). Carbamidomethylation of cysteine was set as a fixed modification; oxidation of methionine, proline and lysine were allowed as variable modifications. Only tryptic peptides were considered, with up to one missed cleavage permitted. Monoisotopic precursor mass values were used, and only +2, +3 and +4 charged precursor ions were considered. Mass tolerances for precursor and fragment ions were 5 ppm and 0.5 Da respectively. MS datasets were validated using rigorous statistical algorithms at both the peptide and protein level implemented in Scaffold 4 (version 4.6; Proteome Software, Portland, OR, USA) [73]. Protein identifications were accepted upon assignment of at least two unique validated peptides with ≥90% probability, resulting in ≥99% probability at the protein level and an estimated 0.1% protein false discovery rate (FDR) for all datasets.

### Proteomic mass spectrometry whole protein data quantification

Relative protein abundance was calculated using peptide intensity by Progenesis LCMS (Non Linear Dynamics Ltd, Newcastle upon Tyne, UK). Orbitrap MS data was loaded and alignment of chromatograms carried out using the automatic alignment algorithm, followed by manual validation and adjustment of the aligned chromatograms. All features with greater than one isotope were used for peptide identifications. Progenesis created the peak list file that was exported and searched in Mascot. Spectra were extracted using extract_msn (Thermo Fisher Scientific, Waltham, MA, USA) executed in Mascot Daemon (version 2.5.1; Matrix Science, London, UK) and imported back into Progenesis to acquire intensity data. Peptide and protein data were then exported from Progenesis as .csv files to be analysed in Excel.

### Hierarchical clustering analysis

Z-transformed mean normalised protein intensities were used for hierarchical clustering of proteomic data and solubility profiling across fractions. Agglomerative hierarchical clustering was performed using MultiExperiment Viewer (version 4.9.0) [74]. Protein hits were hierarchically clustered based on Euclidean distance, and distances between hits were computed using a complete-linkage matrix. Clustering results were visualised using MultiExperiment Viewer (version 4.9.0).

### Peptide identification for peptide location fingerprinting

To interrogate the basement membrane proteins specifically, only the ECM-enriched sample datasets (fraction 3) were analysed by PLF. Raw LC-MS/MS files (.raw) were converted to peak lists files (.mgf) using RawConverter (Scripps Research Institute; CA, USA) [80]. MS/MS peptide ions were then searched with Mascot (v2.5.1, Matrix Science, MA, USA) against the Swissprot and Trembl databases (2018) using the following parameters: taxonomy – *Mus musculus* (84,416 proteins and isoforms); fragment tolerance – 0.5 Da and parent tolerance – 5.0 ppm (both monoisotopic); fixed modification – +57 Da on cysteine (C, Carbamidomethylation); variable modification – +16 Da on lysine (L, oxidation), methionine (M, oxidation) and proline (P, hydroxylation), and +42 Da on the peptide N-terminus (acetylation); enzyme – trypsin; max missed cleavages – 1; peptide charge – 2+, 3+.

Search result files (.dat) were exported from Mascot and loaded into Scaffold 5 (Proteome Software; OR, USA), where proteotypic peptides were thresholded to ≥95% peptide probability, giving a low FDR of 0.7% (calculated by Scaffold 5 using the PeptideProphet algorithm [WA, USA] [73] with peptide probabilities assigned by the Trans Proteomic Pipeline). These high confidence, proteotypic peptides, and their respective spectrum matches were exported (.csv) for PLF analysis.

### Peptide location fingerprinting

Peptide location fingerprinting analysis was performed using our published Manchester PLF webtool (MPLF; https://www.manchesterproteome.manchester.ac.uk/#/MPLF) as previously described [29,30,51,81]. Peptide list files (.csv), exported from Scaffold 5, were uploaded to the MPLF webtool. Only proteins present in both Influenza-resolved and control groups were interrogated (to specifically assess structure-associated differences between groups). Protein primary sequences were bioinformatically divided in 50 aa-sized segments. Peptides were mapped, and exclusive spectrum count were summed, within each segment. Peptides spanning two adjacent segments were counted in both. To remove the skew caused by differences related to whole protein relative abundance and enhance protein regional differences in peptide yield, summed counts per segment were median normalised against protein-specific, experiment-wide total spectrum counts. Normalised, summed counts per protein segment were then averaged per group (Influenza-resolved and control) and statistically tested using unpaired, Bonferroni-corrected, repeated measures ANOVAs to identify significant differences (p ≤ 0.05) along protein structures. Fluctuations of peptide yields along protein structures were visualised by subtracting the average, normalised counts in the control group from that of the Influenza-resolved group and dividing them by the segment length (50 aa), to reveal protein regions with the largest, regional differences.

### Ageing Dataset Collection

The LC-MS/MS data from ECM-enriched, young and aged mouse lung samples was originally sourced as a publicly available dataset from the Proteomics Identification Database (PRIDE; PXD012307), the generation of which has been described previously by Angelidis and Simon *et al* [28]. The PLF analysis of young vs aged groups (which was used here to compare between Influenza-resolution and ageing: **Figure 4**), was previously performed and published by our group [29]. PLF analysis output files (.csv), biomarker candidate shortlists and their peptide yield visualisations are all publicly available for download by visiting our MPLF webtool [81], under the “pre-analysed data section” (Old vs Young Mouse Lung ECM Fraction) found at https://www.manchesterproteome.manchester.ac.uk/#/MPLF. This analysis, and its associated visualisations, were downloaded and used here in the comparison between Influenza-resolution and ageing **(Figure 4).**

### Statistics

Unpaired, Bonferroni-corrected, repeated measures ANOVAs were used to identify significant differences (p ≤ 0.05) along protein structures measured by PLF **(Figures 3, 4)** and in collagen IV and laminin chain mRNA transcripts measured using RT-PCR **(Figure 5)**. Normalised protein abundances were statistically analysed by the Progenesis QI pipeline which uses an unpaired repeated measured ANOVA **(Tables 1, 2)**.

## Data Availability

PLF analysis output files (.csv), biomarker candidate shortlists and their peptide yield visualisations are all available under the “Location Fingerprinter” tab on our open access MPLF webtool [81] at https://www.manchesterproteome.manchester.ac.uk/#/MPLF, [25,27]. For peer review, please use the login – username: Reviewer ; password: Proteome2020 . The mass spectrometry proteomics data have been deposited to the ProteomeXchange Consortium via the PRIDE partner repository with the dataset identifier PXD054592 and 10.6019/PXD054592. For peer review this can be accessed via the reviewer account: Username; Password: Y4MfBVvU8Y8q. PLF and mass spectrometry data will be made public upon journal acceptance.

## Acknowledgements

We would like to thank the University of Manchester Biological Mass Spectrometry (BioMS; RRID: SCR_020987) facility for their help in generating and interpreting the MS data, and IT services and Research IT for their aid in hosting, running and maintenance of the MPLF webtool. LC, SK and TH are supported by the Wellcome Trust (202865/Z/16/Z).

## Author Contributions

Conceptualization, TH, AE, RL; methodology, OB, AE, RL; software, AE, MO; formal analysis, OB, TH, AE; investigation, OB, TH, AE; resources, TH; data curation, BO, AE writing—original draft preparation, TH, AE; writing—review and editing, OB, MO, SK, CJ, RH, TH, AE; visualization, OB, TH, AE; supervision, TH, AE; project administration, TH, AE; funding acquisition, TH.

