## Supplementary material for "Lung basement membranes are compositionally and structurally altered following resolution of acute inflammation": Figure S1

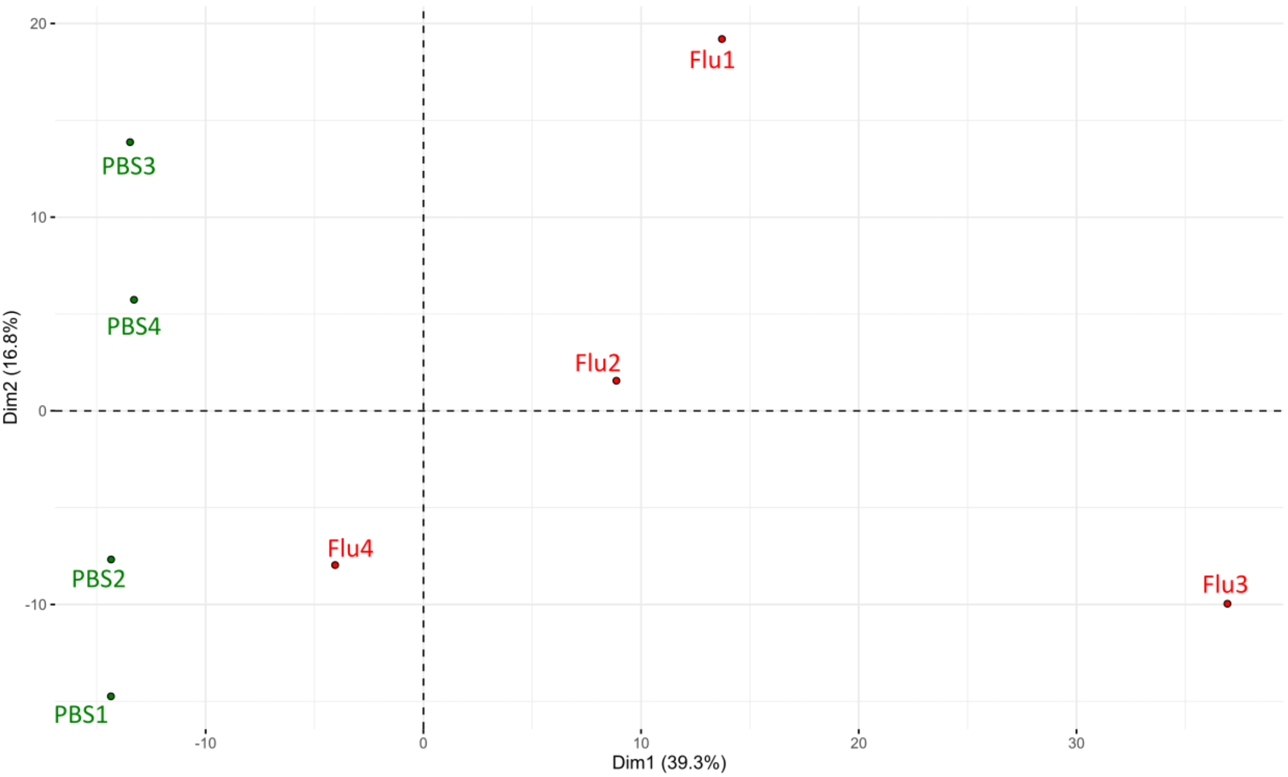


**Figure S1.** Principal component analysis (performed in R using *FactoMineR*; visualised using *factoextra*) of peptide spectral counts identified from flu-resolved and control mouse lung samples, used for PLF. Control (green) and post-flu (red) sample groups displayed good data separation along PC1.
